# High-sensitivity vision restoration via ectopic expression of chimeric rhodopsin in mice

**DOI:** 10.1101/2020.09.17.301523

**Authors:** Yusaku Katada, Kazuho Yoshida, Naho Serizawa, Kenta Kobayashi, Kazuno Neghisi, Hideyuki Okano, Hideki Kandori, Kazuo Tsubota, Toshihide Kurihara

**Affiliations:** Laboratory of Photobiology, Keio University School of Medicine, Shinanomachi, Shinjuku-ku, Tokyo, 160-8582, Japan; Department of Ophthalmology, Keio University School of Medicine, Shinanomachi, Shinjuku-ku, Tokyo, 160-8582, Japan; Department of Life Science and Applied Chemistry, Nagoya Institute of Technology, Nagoya, Aichi, 466-0061, Japan; Section of Viral Vector Development, Center for Genetic Analysis of Behavior, National Institute for Physiological Sciences, National Institutes of Natural Sciences, Okazaki, Aichi, 444-8585, Japan; Department of Physiology, Keio University School of Medicine, Shinanomachi, Shinjuku-ku, Tokyo, 160-8582, Japan; Tsubota Laboratory, Inc., Tokyo, 160-0016, Japan

## Abstract

Photoreception requires amplification by mammalian rhodopsin through G protein activation, which requires a visual cycle. To achieve this in retinal gene therapy, we incorporated human rhodopsin cytoplasmic loops into *Gloeobacter* rhodopsin, thereby generating *Gloeobacter* and human chimeric rhodopsin (GHCR). In a murine model of inherited retinal degeneration, we induced retinal GHCR expression by intravitreal injection of a recombinant adeno-associated virus vector. Retinal explant and visual thalamus electrophysiological recordings, behavioral tests, and histological analysis showed that GHCR restored dim-environment vision and prevented the progression of retinal degeneration. Thus, GHCR may be a potent clinical tool for the treatment of retinal disorders.

**One Sentence Summary:** Optogenetic therapy with Gloeobacter and human chimeric rhodopsin resulted in highly sensitive visual restoration and protection effects.

## INTRODUCTION

Inherited retinal degeneration (IRD) is a major cause of vision loss. More than 2 million people worldwide are blind due to IRD*(1)*, and few effective treatments exist. For retinitis pigmentosa (RP), one of the most common forms of IRD, previous studies have reported vision restoration in animal models using various molecules as optogenetic actuators*(2–9)*. In addition, clinical trials are under way to investigate the effects of introducing channelrhodopsin 2 (RST-001, ClinicalTrials.gov Identifier: NCT01648452) and ChrimsonR (GS-030, ClinicalTrials.gov Identifier: NCT03326336) into retinal ganglion cells (RGCs) via gene transduction achieved by intravitreal injection of recombinant adeno-associated virus (rAAV). The first clinical case report on optogenetic therapy was recently reported*(10)*. However, microbial opsins, such as channelrhodopsin 2, require high light intensity, such as outdoor light intensity levels, to function*(11–13)*. They cannot restore vision in dimly lit environments, such as indoors or at night, and strong light irradiation can promote retinal degeneration*(14, 15)*. Physiological photoreception mediated by mammalian rhodopsin, however, relies on amplification through G protein activation. Although the introduction of vertebrate opsin improved photosensitivity in mice*(9, 16)*, it is unclear how the chromophore retinal is metabolized in the retina where the visual cycle is broken. Animal rhodopsin also causes toxicity if all-trans retinal is not properly metabolized*(17, 18)*, and is, thus, hampered by safety and stability concerns in terms of clinical application.

Because of the above limitations of animal visual opsins, one attempt to circumvent them is the chimeric rhodopsin of melanopsin and G protein-coupled receptor (GPCR)*(8, 19)*. Melanopsin is a non-visual opsin, and despite being an animal opsin, it is not easily photobleached. However, it has a “bistable” photo-cycle and requires different wavelengths of light for conformational change, which may result in unnatural appearance*(20, 21)*.

Therefore, a chimeric rhodopsin of microbial opsin and GPCR*(22–24)*, is not photo-bleached and is a monostable pigment like visual opsin, but may be able to achieve highly sensitive visual restoration via G protein stimulation.

In this study, to achieve light sensitivity, stability, and safety, we attempted to restore vision in mice using *Gloeobacter* and human chimeric rhodopsin (GHCR)*(23, 24)*.

## RESULTS

### Design of GHCR

Although there is no sequence identity between microbial and animal opsin, both possess similar chromophore (retinal) and protein (seven-transmembrane helix) structures. As we previously reported*(24)*, to generate GHCR, we replaced the second and third intracellular loops of *Gloeobacter* rhodopsin with human sequences and introduced the E132Q mutation (**Figure S1**). Previous work has shown that GHCR induces G protein activation *in vitro(24)*.

### Restoring light-evoked activity in the retina with GHCR

We injected a viral vector (rAAV-DJ or rAAV-2) containing the GHCR coding sequence under the control of the hybrid promoter comprising the CMV immediate-early enhancer, CBA promoter, and CBA intron 1/exon 1, known as the CAGGS promoter, (CAGGS-GHCR; **Figure 1a**) into the vitreous humor of 10-week-old *rd1* mice. We adopted the rAAV-DJ vector to achieve more efficient, widespread gene transfer*(25, 26)*, and used rAAV-2 as a benchmark, as it has already been used in the clinic*(27)*. The retinas were harvested 2–4 months later. Enhanced green fluorescent protein (EGFP) reporter gene expression was observed in the retina and in both the ganglion cell layer and the inner nuclear layer (**Figure S2a, b**). To evaluate the function of ectopically expressed GHCR in the mouse retina, we performed multi-electrode array (MEA) recording to record the extracellular potential of RGCs (**Figure S2c**). As a result of photoreceptor degeneration, the untreated control retina showed no RGC response as detected by MEA (**Figure 1b**). In contrast, the treated retinas showed obvious light-induced responses down to 10^14^ photons/cm^2^/s of white light-emitting diode (LED) irradiation (**Figure 1c**).

**Figure 1.**
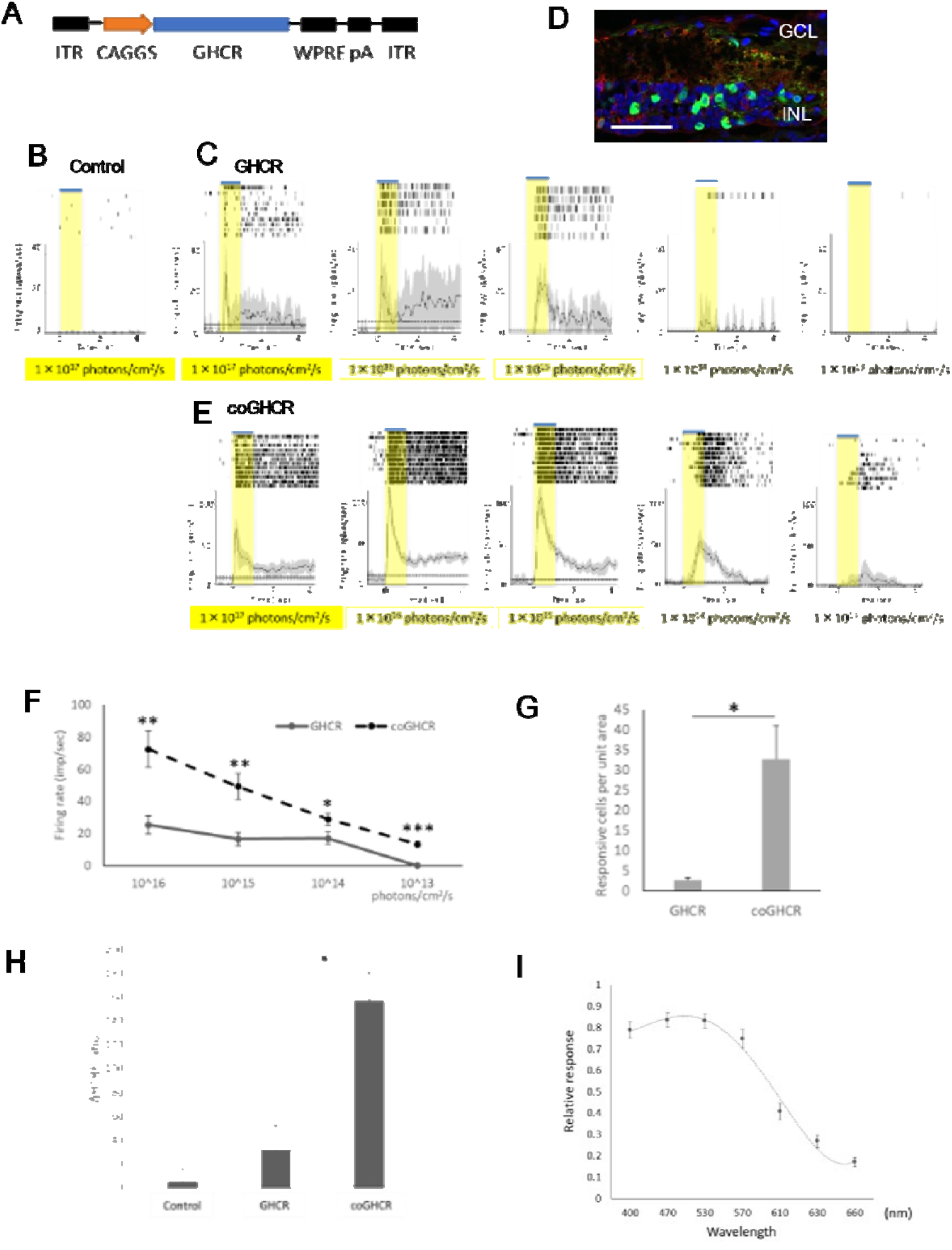
Ectopic GHCR expression restores light responses in the *rd1* mouse retina. (a) DNA expression cassette schematic. The GHCR coding sequence is driven by the CAGGS promoter, flanked by inverted terminal repeats (ITR), and stabilized by a polyadenylation signal sequence (pA) and a woodchuck hepatitis posttranscriptional regulatory element (WPRE). (b, c, e) Raster plots and peri-stimulus time histograms for light stimulation of control (AAV-DJ-CAGGS-EGFP) (b), GHCR-treated (AAV-DJ-CAGGS-GHCR) (c), and coGHCR-treated (AAV-DJ-CAGGS-coGHCR) mice (e). Responses to exposure to a white LED with varying light intensity for 1.0 s. Gray shading around the averaged traces represents the standard error of the mean (SEM). (d) Confocal image of a transverse *rd1* mouse retina section 2 months after AAV-DJ-CAGGS-coGHCR intravitreal injection. Green, FLAG tag antibody signal (vector); red, PKCα signal (bipolar cells); blue, 4′,6-diamidino-2-phenylindolenuclear (DAPI) counterstaining. Scale bar, 50 μm. (f) Quantitation of the firing rates of RGCs transduced with GHCR or coGHCR at the indicated light intensity. (g) Histogram showing the number of RGCs that responded to light per unit area (2.6 mm^2^) of the retinas of GHCR- or coGHCR-treated mice (n = 3 each). (h) Changes in cAMP consumption in response to Gi/o-coupled G-protein-coupled receptor activation in HEK293T cells transfected with GHCR and coGHCR (n = 3 each). (i) Spectral sensitivity induced by coGHCR (n = 23 cells each). Error bars represent the SEM. Data were analyzed with Student’s two-tailed t-test in (f, g) and one-way analysis of variance (ANOVA) and Tukey’s multiple comparison test in (h); * represents p ≤ 0.05, ** represents p ≤ 0.01, and *** represents p ≤ 0.001. GCL, ganglion cell layer; INL, inner nuclear layer.

Next, to create a stable vector for human gene therapy, we designed a codon-optimized version of GHCR (coGHCR) and fused the ER2 endoplasmic reticulum (ER) export signal to its C-terminus to increase gene expression levels. Immunolabelling revealed expression across the whole retina, including in the bipolar cells, of treated *rd1* mice (**Figures 1d**). As a result, the firing rate increased significantly, and a photoresponse was confirmed down to 10^13^ photons/cm^2^/s, which had not observed before optimization (**Figure 1e, f**). The retinas of WT mice were highly responsive to all light stimulus levels under dark-adapted conditions, but under light-adapted conditions, the firing rate was also modulated in response to light stimulus intensity, and coGHCR response was similar to the light-adapted conditions in WT mice (**Figure S2e**). No photoresponse to any light stimulus level was obtained from control untreated mice. Moreover, the number of firing cells per unit area also increased significantly (**Figure 1g**). Since rhodopsin shows selectivity for Gi/o class G proteins upon heterologous expression*(28–31)*, we measured Gi/o activation with a homogeneous time-resolved fluorescence (HTRF) cyclic adenosine monophosphate (cAMP) assay. We observed a 5-fold increase in activation in coGHCR-treated compared with GHCR-treated mice (**Figure 1h**). The maximum spectral sensitivity of retinas treated with coGHCR was around 500 nm, and a photoresponse was obtained even upon stimulation with light with a wavelength >600 nm (**Figure 1i**).

### Restoration of visual cortex responses by GHCR

To investigate whether retinal light responses were transmitted to the visual cortex, we then examined visual evoked potentials (VEPs) generated by the visual cortex (**Figure 2a**). The output from the RGCs is sent through their axons (optic nerve) to the lateral geniculate nucleus (LGN) of the thalamus, which is a region of the diencephalon, then from the LGN to the primary visual cortex in the occipital lobe of the cerebral cortex. For these experiments, we used *rd1* mice in which both eyes had been treated with the AAV-DJ-CAGGS-GHCR, AAV-DJ-CAGGS-coGHCR, or control EGFP (AAV-DJ-CAGGS-EGFP) vectors. Significant VEPs were not detected in the control or GHCR-treated mice. In contrast, VEPs were observed in coGHCR-treated mice (**Figure 2b**). In response to 3 cd·s/m^2^ light stimulation, the average VEP amplitude in coGHCR-treated mice was significantly higher (56.4 μV; n = 6) than those in GHCR-treated mice (22.1 μV; n = 8) and control mice (17.9 μV; n = 6) (**Figure 2c**). Based on this result, all subsequent experiments were performed using coGHCR.

**Figure 2.**
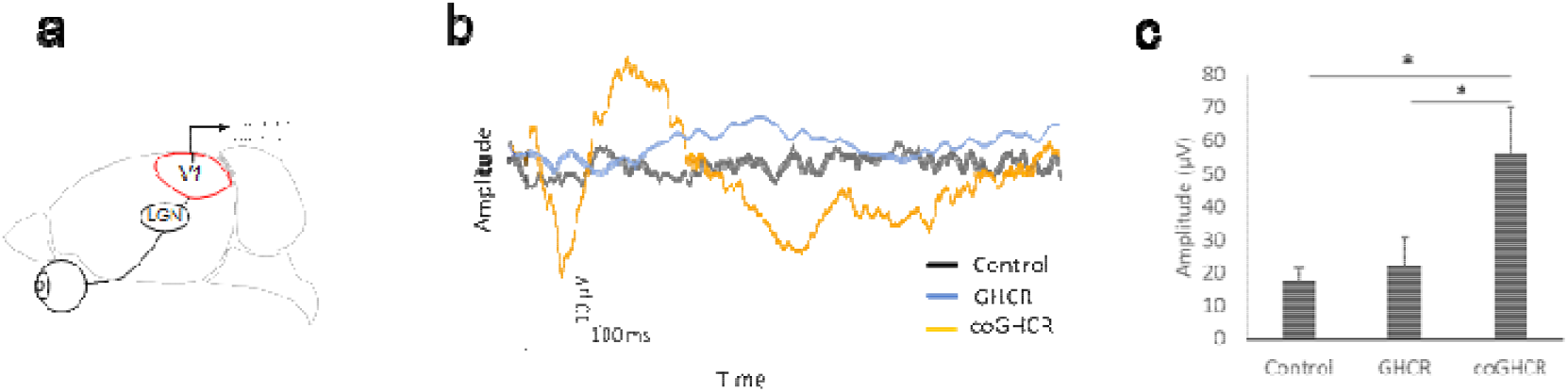
coGHCR restored vision in *rd1* mice through the primary visual cortex. (a) Schematic view of the VEP recording strategy. (b) Representative VEP traces from GHCR-treated, coGHCR-treated, and control mice. (c) The average amplitude of the VEPs in the control (AAV-DJ-CAGGS-EGFP, n = 6), GHCR-treated (AAV-DJ-CAGGS-GHCR, n = 8), and coGHCR-treated (AAV-DJ-CAGGS-coGHCR, n = 6) mice. The stimulus was a white LED flash (3 cd·s/m^2^). Signals were low-pass filtered at 300 Hz and averaged over 60 trials. Error bars represent the SEM. Data were analyzed with one-way ANOVA and Tukey’s multiple comparison test; * represents p ≤ 0.05. V1, visual cortex.

### Characterization of the in vivo responses restored by GHCR transduction

Next, light-dark transition (LDT) testing was performed to investigate whether ectopic expression of coGHCR in degenerating retinas led to behavioral changes due to vision restoration (**Figure 3a**). Rodents with intact vision tend to stay in dark places as they are nocturnal and feel uneasy in bright environments, whereas blind rodents spend roughly half of their time in bright places. The coGHCR-treated mice spent significantly less time in the bright area compared with the untreated *rd1* mutant mice (**Figure 3b**), thereby confirming vision restoration via behavioral analysis. And the visual restoration effect was still maintained after two years (**Figure 3c**). Furthermore, in order to directly compare the effects of coGHCR with genes in clinical trials, we treated *rd1* mice with chimeric rhodopsin (AAV-6-CAGGS-coGHCR), microbial opsin (AAV-6-CAGGS-ChrimsonR*(32)*), animal rhodopsin (AAV-6-CAGGS-human rhodopsin), or the control EGFP (AAV-6-CAGGS-EGFP) vector. At an illuminance of 3,000 lux, a significant reduction in the time spent in the bright half of the observation area was noted for coGHCR-treated mice (0.32; n = 6) compared with control mice (0.50; n = 8) (**Figure 3d**). A similar tendency was observed in ChrimsonR-treated mice (0.36; n = 6). However, no obvious change was observed in human rhodopsin-treated mice (0.48; n = 6). When the experiment was carried out at an illumination of 10 lux, human rhodopsin-treated mice showed a significant change in the time spent in the bright area (0.40; n = 6), whereas ChrimsonR-treated mice did not show an obvious change (0.55; n = 6) (**Figure 3e**). The coGHCR-treated mice again spent significantly less time in the bright area illuminated at 10 lux (0.40; n = 6).

**Figure 3.**
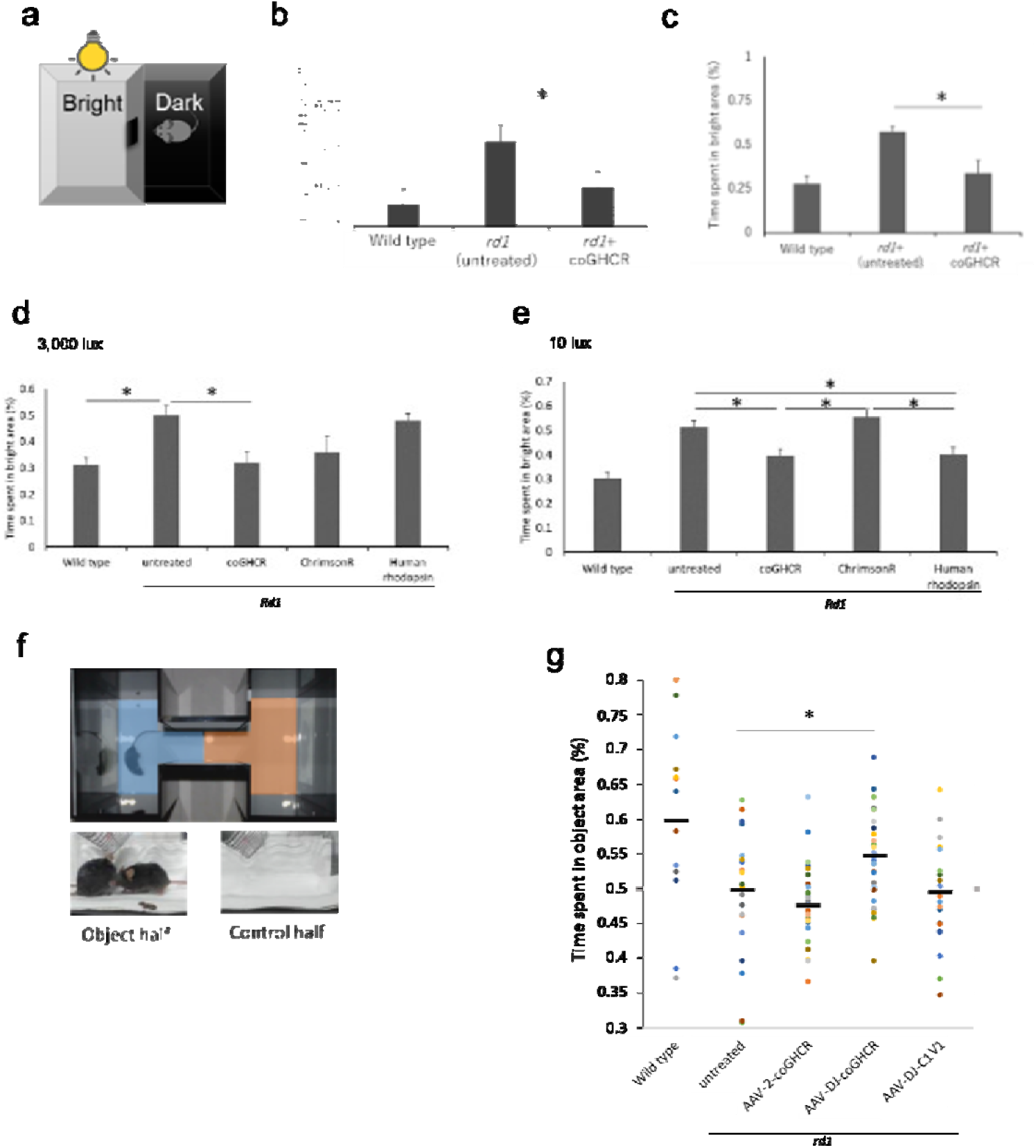
coGHCR-treated mouse behavior indicated vision restoration. (a) LDT testing schematic. Mice were tested in a 30 × 45 × 30-cm box with equally sized bright and dark chambers connected by a 5 × 5-cm opening, across which the mice could move freely. (b, c) Percentage of time spent in the bright area (total, 10 min) by wild type (n = 4), and control (AAV-DJ-CAGGS-EGFP) (n = 7 in (b) and n = 4 in (c)) and coGHCR-treated (AAV-DJ-CAGGS-coGHCR) *rd1* mice (n = 6). LDT test at 3 months (b) and 2 years (c) after treatment, 10 lux illumination. (d, e) The percentage of time spent in the bright area (total, 10 min) by wild type (n = 6), and control (AAV-6-CAGGS-EGFP) (n = 8), coGHCR-treated (AAV-6-CAGGS-coGHCR) (n = 6), ChrimsonR-treated (AAV-6-CAGGS-ChrimsonR) (n = 6), and human rhodopsin-treated (AAV-6-CAGGS-human-rhodopsin) *rd1* mice (n = 6). LDT test with 3,000 lux (d) and 10 lux (e) illumination. (f) VRT setup. Time spent in areas showing a video of mice fighting (object half, blue) or an empty cage (control half, red) was measured. (g) Distribution of time spent in the object half by wild type (n = 14), and control (no treatment) (n = 23), AAV-2-coGHCR-treated (AAV-2-CAGGS-coGHCR) (n = 30), AAV-DJ-coGHCR-treated (AAV-DJ-CAGGS- coGHCR) (n = 33), and AAV-DJ-C1V1-treated (AAV-DJ-CAGGS-C1V1) *rd1* mice (n = 20). LDT test with 10 lux (d) and 3,000 lux (e) illumination. Black line, average value. Error bars represent the SEM. Data were analyzed with one-way ANOVA and Tukey’s multiple comparison test; * represents p ≤ 0.05.

### Restored object recognition function upon GHCR gene therapy

LDT testing measures only light and dark discrimination. Visual recognition testing (VRT) was performed to evaluate whether the mice could recognize an object with the restored level of vision. Mice use vision for their cognitive functions, and are attracted to fighting videos*(33–35)*. We examined mice in a place preference apparatus with a tablet showing a fighting video (**Figure 3f**). The ratio of the time spent in the area with the fighting compared with the time spent in the control area (showing a video of an empty cage with the same illuminance) over 15 minutes was measured. The coGHCR-treated (AAV-DJ-CAGGS-coGHCR) mice spent significantly more time in the fighting video half of the apparatus (0.55, n = 33) than the untreated *rd1* mice (0.50, n = 30). On the other hand, microbial opsin-treated (AAV-DJ-CAGGS-C1V1*(36)*) mice spent roughly equivalent time in each half (0.49, n = 20) (**Figure 3g**).

### GHCR protective effects against retinal degeneration

We employed another mouse model of retinal degeneration using mice with the P23H *RHO* mutation, referred to as P23H mice*(37)*. P23H mice were selected to evaluate the protective effect because they have slower retinal degeneration than *rd1* mice. We subretinally delivered AAV DJ-CAGGS-coGHCR and the control (AAV DJ-CAGGS-EGFP) vector into postnatal day (PND) 0–1 mouse retinas, targeting the outer retina, and quantified the protective effects of the vector via morphological and electrophysiological examination. Subretinal injection of AAV-DJ efficiently induced gene expression in the murine outer retina (**Figure 4a**). Optical coherence tomography (OCT) showed that the outer retinal thickness (ORT), which is the thickness from the outer nuclear layer (ONL) to the rod outer segment (ROS), of coGHCR-treated mice (50.0 μm; n = 13) was significantly greater than that of the control mice (42.7 μm; n = 10) at PND 30 (**Figure 4b, c**). The ORT of the treated mice remained significantly greater than that of control mice until PND 50 (**Figure S3**).

**Figure 4.**
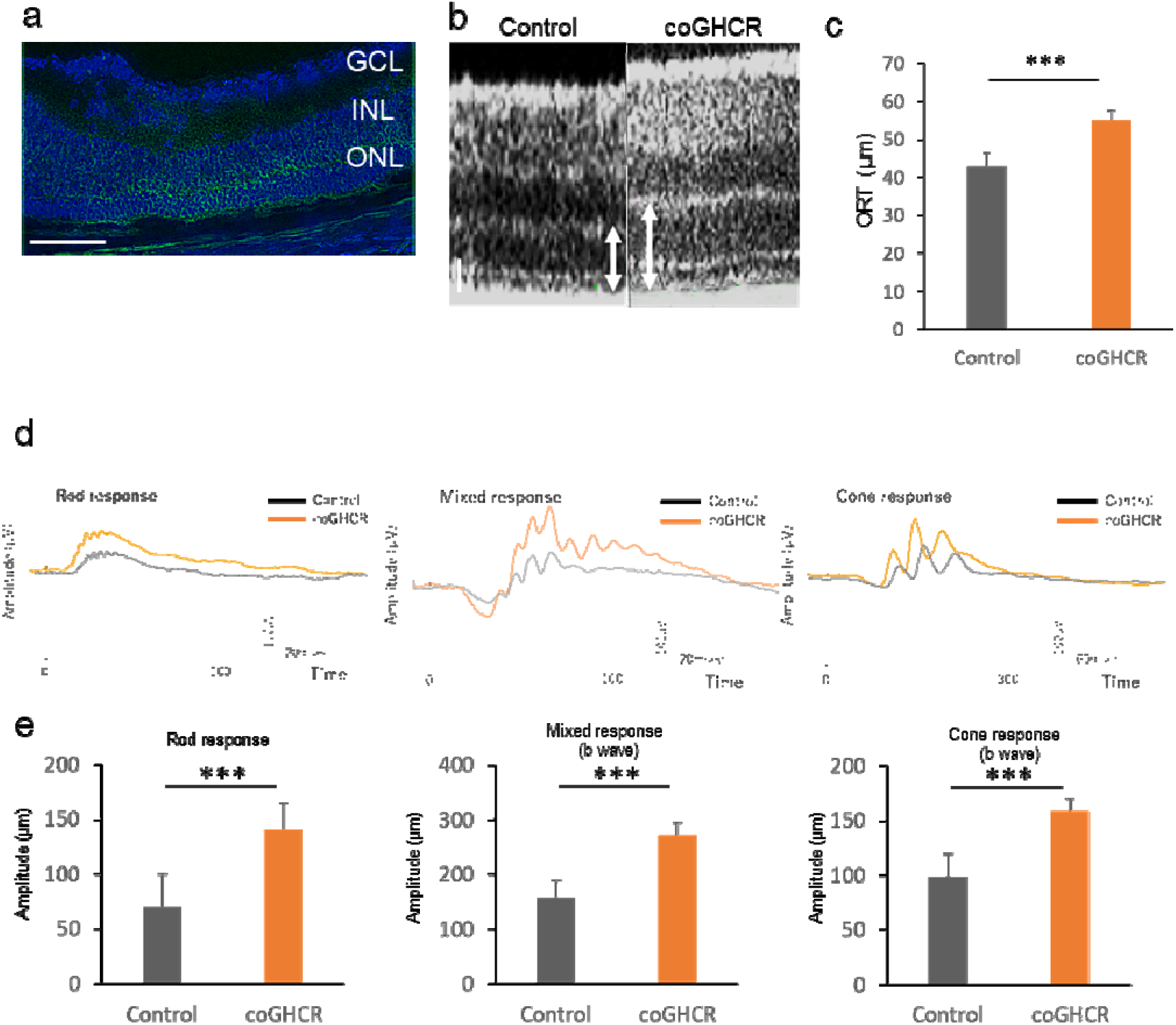
Ectopic coGHCR expression protects against photoreceptor degeneration. (a) Confocal image of a transverse section through the P23H retina 2 months after AAV-DJ-CAGGS-coGHCR subretinal injection. Green, FLAG tag fused to the C-terminus of coGHCR; blue, DAPI nuclear counterstaining. Scale bar, 100 μm. (b) OCT retinal image sections from coGHCR-treated and control (AAV-DJ-CAGGS-EGFP subretinally injected) mice at PND 30. The white arrow indicates the measured ORT (from ONL to cone outer segment). Scale bar, 20 μm. (c) Histogram of the measured ORT of the coGHCR-treated (n = 13) and control mice (n = 10) at PND 30. (d, e) Representative ERG waveforms (rod response, mixed response, and cone response) of coGHCR-treated (n = 14) and control mice (n = 9) (d). Histograms of the average ERG amplitudes from panel d at PND 30 (e). Error bars represent SEM. Data were analyzed with the unpaired t-test; *** represents p ≤ 0.001. GCL, ganglion cell layer; INL, inner nuclear layer.

Electroretinography (ERG) revealed that the treated mice had larger rod, mixed, and cone response amplitudes (141.2 μV, 271.4 μV, and 159.0 μV, respectively; n = 9) than the control mice (70.4 μV, 158.7 μV, and 99.1 μV, respectively; n = 14) at PND 30 (**Figure 4d, e**). All amplitudes in the control mice gradually decreased, whereas all amplitudes in the coGHCR-treated mice continued to increase until PND 42 (**Figure S4a, b c**). Thereafter, the amplitudes in the treated mice also gradually decreased, although they remained significantly higher than those in the control mice until PND 66.

We also performed terminal deoxynucleotidyl transferase dUTP nick end labeling (TUNEL) to detect apoptosis in the retinas. The number of TUNEL-positive cells in the coGHCR-treated mouse ONL (289.7 cells; n = 3) was significantly lower than that in the control mouse ONL (67.3 cells; n = 3) at PND 31 (**Figure 5a–c**).

**Figure 5.**
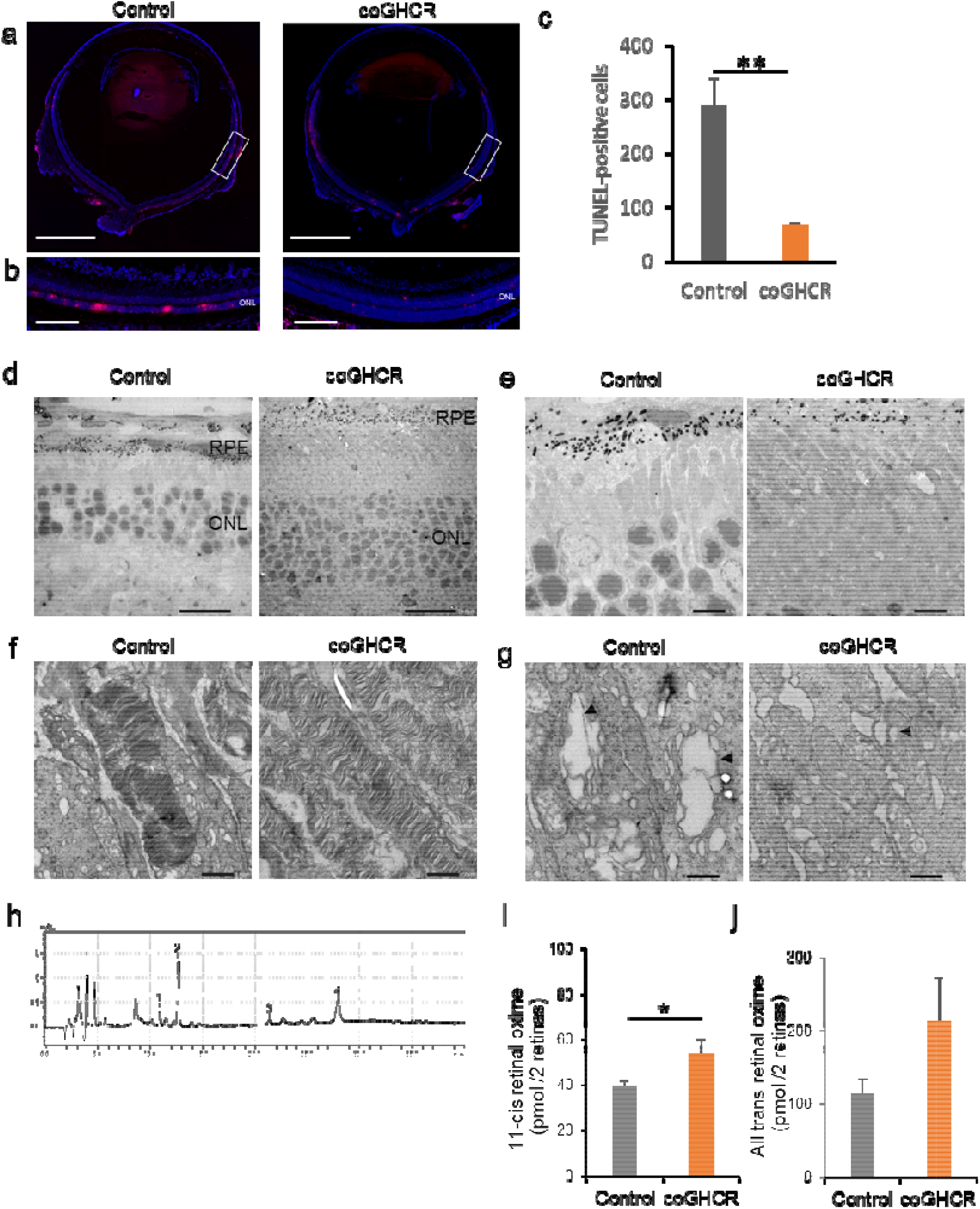
coGHCR treatment suppressed retinal apoptosis and ER stress. (a, b) TUNEL-stained transverse sections (a) and enlarged images of the white squares (b) of coGHCR-treated and control (AAV-DJ-CAGGS-EGFP subretinally injected) mouse retinas at PND 31. Red, TUNEL-positive cells; blue, DAPI nuclear counterstaining. Scale bar, 1,000 μm in (a) and 100 μm in (b). (c) Histogram of the number of TUNEL-positive cells in the ONLs of coGHCR-treated (n = 3) and control mice (n = 3) at PND 31. (d) TEM images of transverse sections from coGHCR-treated and control mice at PND 31, showing the outer retinal layer (d), the outer segment at low magnification (e) and high magnification (f), and the inner segment (g). The arrowhead indicates swollen ER. Scale bar, 20 μm in (d), 5 μm in (e), 1 μm in (f), and 500 nm in (g). (h) Chromatograms of retinal in mouse retina analyzed by HPLC. 15 h dark adapted mice were exposed to light of 1000 lux for 10 min and each retina was processed and retinal oximes extracted under dim red light. Peak identification was determined using retinal standard reagents as follows: 1, syn-11-cis-retinal oxime; 2, syn-all-trans-retinal oxime; 3, anti-11-cis-retinal oxime; 4, anti-all-trans-retinal oxime. (i, j) Histogram quantifying the amount of retinal oximes from coGHCR-treated (n = 9) and control mice (n = 9) obtained from HPLC. Error bars represent SEM. Data were analyzed with the unpaired t-test; * represents p ≤ 0.05, ** represents p ≤ 0.01.

To expand these observations, we obtained transmission electron microscopy (TEM) images of transverse sections from PND 31 mice. Consistent with the OCT results, the ONL (**Figure 5d**) and ROS (**Figure 5e**) of coGHCR-treated mice were relatively intact compared with those of controls, and the ROS structure was less disorganized (**Figure 5f**). In addition, coGHCR-treated mice had less swelling of their ER, a feature that is indicative of ER stress (**Figure 5g**).

We also performed western blotting to investigate ER stress. Expression of the ER stress marker ATF4 was significantly lower, and expression of BiP, PERK, ATF6, and pIRE1 tended to be lower in treated mice than in control mice at PND 14 (**Figure S5a, b**).

Since retinoid levels are known to affect ER stress and retinal degeneration, retinoid analysis of the treated eyes was performed. The amount of retinal was measured by HPLC using the retinal oxime method after 10 minutes of exposure to 1000 lux, a fluorescent lighting level assuming a normal indoor environment. The results showed that 11-cis retinal oximes was significantly elevated in the treated eyes (54.1 ± 18.2 pmol/ 2 retinas; n = 9) versus controls (39.5 ± 6.5 pmol/2 retinas; n = 9) (**Figure 5h, i**). No obvious changes in the amount of all-trans-retinal oxime were observed (**Figure 5j**).

## DISCUSSION

Because the phenotype of retinal degeneration is common across cases of retinitis pigmentosa, regardless of genotype, the strategy of optogenetic therapy has great potential as a universal therapeutic approach. It aims to target non-photoreceptive surviving neurons in the retina, such as retinal ganglion cells and bipolar cells, and convert them to photoreceptive.

In this study, we demonstrated that ectopic expression of coGHCR is an effective method of optogenetic vision restoration in mice with retinal degeneration. MEA revealed that photoresponses were maintained for retinal irradiance levels as low as 10^13^ photons/cm^2^/s. This is consistent with the response of the treated mice to 10 lux illumination in the behavioral test, and represents a significant improvement in sensitivity compared with that observed in previous studies of vision restoration with microbial opsins (threshold: 10^14^ to 10^17^ photons/cm^2^/s)*(2–7)*, LiGluR/MAG photoswitches (threshold: 10^15^–10^16^ photons/cm^2^/s)*(38, 39)*, or photoactivated ligands (AAQ threshold: 10^15^ photons/cm^2^/s*(40)* and DENAQ threshold: 4 × 10^13^ photons/cm^2^/s*(41)*). Although some vectors restored greater sensitivity, such as human rhodopsin*(9)*, cone opsin*(16)* and Opto-mgluR6 (10^12^ photons/cm^2^/s)*(8)*, our LDT results at 3,000 lux (similar to a cloudy outdoor environment) suggest that photobleaching of rhodopsin like these does not work in bright environments. coGHCR is adaptable to a light environment ranging from at least 10 lux (similar to a night light levels with streetlights) to 3,000 lux, and is, thus, a suitable single-opsin vision restoration tool.

Furthermore, the typical channelrhodopsins have a spectrum limited to blue light*(42)*, which limits their use as a visual restoration tool. On the other hand, GHCR has a spectrum peak around 500 nm and facilitates responses to red light. Irradiation of high-energy light such as blue light can cause phototoxicity and cell death due to generation of free radicals*(43)*. Therefore, there are concerns about phototoxicity in optogenetic tools that operate under blue light, such as channelrhodopsin, and long wavelength-shifted opsins have been developed*(44)*. In this regard, the GHCR has the advantage of being highly sensitive and having a peak at intermediate (green) wavelengths, making it responsive to short and long wavelengths and less likely to exceed safe limits of light intensity*(45)*. In addition, behavioral tests showed that coGHCR enabled responses to both sustained and transient stimulation lasting 10 ms. These findings suggested that coGHCR gene therapy can restore sensitivity to multiple light environments encountered in daily life.

The ERG amplitudes in coGHCR-treated mice continued to increase until PND 42, likely because the coGHCR-mediated signal was additive with the innate amplitude. This is consistent with the fact that gene expression of the AAV-DJ vector peaks at approximately 1.5 months after administration*(25)*. We observed no apparent changes in the shapes of the ERG waveforms in the coGHCR-treated mice. The visual restoration effect was also maintained for two years, which shows promise for long-term pharmacological effects and safety.

coGHCR has Gt activity derived from rhodopsin*(24)*. Gt is also known to be cross-linked with Gi/o*(46)*, and this was confirmed (**Figure 1h**). Although this study used a ubiquitous promoter, which cannot be fully confirmed, Gi/o is generally expressed specifically in ON-type bipolar cells*(47, 48)*, where the light-responsive signal is likely to have been generated. When coGHCR is expressed ectopically in ON bipolar cells, it is expected to inhibit responses. However, the restored responses observed by MEA were all ON responses. In addition, the electrophysiological and behavioral results were similar to physiological responses, and no reversal reaction observed. In *rd1* mice, photoreceptors are mostly lost by 4 weeks after birth and no optical response is obtained after 7 weeks at the latest*(49, 50)*. Therefore, responses from residual photoreceptors are unlikely in this study. A similar phenomenon has been confirmed in previous studies; the excitatory response is hypothesized to result from disinhibition of inhibitory amacrine cells*(6, 8, 9)*.

The safety of ectopic expression of opsins, such as channelrhodopsin 2, has been previously reported*(3, 51, 52)*. To our knowledge, this is the first report of their protective effects against retinal degeneration. In vitro studies have shown that the P23H opsin is misfolded and retained in the ER*(53)*. ER retention of P23H opsin can induce the unfolded protein response, leading to apoptosis*(54–57)*. Our results suggest that expression of coGHCR in the retinal outer layer suppressed ER stress and photoreceptor apoptosis, which led to protection against degeneration. The lack of 11-cis-retinal induces cytotoxicity during the development of ROS in P23H mice*(58)*. In fact, the amount of cis-retinal in the retina was significantly elevated after coGHCR treatment. Since coGHCR uses all-trans retinal as a chromophore, like microbial opsin, it does not consume cis-retinal and is free from photobleaching. Therefore, the expressed coGHCR may suppress cis-retinal consumption via photoreceptor substitution. If this hypothesis is correct, the protection effect of coGHCR may not be applicable to patients with all IRD genotypes. However, there are more than 140 known RP-linked rhodopsin mutations, and those that result in protein misfolding and retention in the ER are the most prevalent*(59, 60)*.

In summary, the coGHCR vector has the advantages of both animal and microbial opsin as a vision regeneration tool. It restores sensitivity and an action spectrum that enables vision in lighting ranging from levels found outdoors to those in dimly lit indoor environments via G protein stimulation without the risk of bleaching; it can also be expected to protect against the progression of retinal degeneration in the majority of IRD patients. These results suggest that coGHCR is worthy of consideration for clinical application as a gene therapy for IRD.

## MATERIALS AND METHODS

Study approval: All of the animal experiments were conducted in accordance with protocols approved by Institutional Animal Care and Use Committee of Keio University School of Medicine (#2808).

Please see supplemental information for detail.

## Author Contribution

Y.K. and T.K. designed the research, wrote the manuscript. Y.K. performed the retinal histology, MEA, ERG and VEP recordings, and WB, TEM, LDT and VRT experiments. N.S. performed HPLC. K.Y. performed plasmid construction. K.K. performed AAV production. Y.K. performed data processing and analysis. K.N., H.K., H.O and T.K. made critical revisions of the manuscript. T.K. supervised the research.

## Acknowledgements

Y.K. was supported by grants from the Keio University Doctorate Student Grant-in-Aid Program and Grant-in-Aid for Research Activity Start-up. T.K. was supported by Grants-in-Aid from Takeda Science Foundation and the Keio University Medical Science Fund. We would like to thank Editage (www.editage.com) for English language editing.

## Declaration of Interests

Y.K., H.K., K.T., and T.K. are inventors on pending patents (PCT/JP2017/031579、 PCT/JP2019/ 1565) related to this work. Y.K., K.T., and T.K. are equity holders in Restore Vision, Inc.

## Notes

### Summary of Updates

Updated supplemental files from docx to PDF again.

