## Supplemental Information for "High-sensitivity vision restoration via ectopic expression of chimeric rhodopsin in mice"

Katada Y et al.

### Supplemental Methods

#### Animals

Mice homozygous for the retinal degeneration alleles $Pde6b^{rd1}$ (C3H/HeJJcl, *rd1*) and WT C57BL/6J were obtained from CLEA Japan, Inc. Mice heterozygous for the retinal degeneration alleles $Rho^{P23H/+}$ （B6.129S6(Cg)-$Rho^{tm1.1Kpal}$/J, P23H） were obtained from Jackson Laboratory. Animals were maintained under 12-h light:12-h-dark conditions. For animals bred in house, littermates of the same sex (male) were randomized to experimental groups. All of the animal experiments were conducted in accordance with protocols approved by Institutional Animal Care and Use Committee of Keio University School of Medicine.

#### Method Details

##### Immunohistochemistry

The protocol for immunohistochemistry was previously described[1]. The retinas were incubated in PBS with 1% Triton X-100 and 0.5% Tween 20 for 1 h at room temperature and in 4% BSA for 1 h at room temperature and then incubated overnight at 4 ˚C with primary antibodies: anti-FLAG (1:500, Merck, Darmstadt, Germany) and anti-PKCα (1:100, Abcam, Cambridge, UK) in blocking buffer. Secondary anti-rabbit, conjugated with Alexa TM488 or 594 (1:1000; Abcam), were applied for 1 h at room temperature.

**Vector production and purification**

GHCR construct were designed as previously reported[2]. GHCR, coGHCR, ChrimsonR, C1V1 amd human rhodopsin genes were cloned to pAAV-CAGGS-MCS. Type 2, 6, DJ serotypes of rAAV vectors were prepared using the AAV Helper Free Packaging System (Cell Biolabs, San Diego, CA, USA). The serotypes were produced in HEK 293T cells using a helper virus-free system and were purified using two $CsCl_2$ density gradients and titrated by quantitative polymerase chain reaction. Final preparations were dialyzed against phosphate-buffered saline (PBS) and stored at -80°C.

**Virus injection**

The mice were anesthetized with a combination of midazolam, medetomidine and butorphanol tartrate at doses of 4 mg/kg, 0.75 mg/kg and 5 mg/kg of body weight and placed on a heating pad that maintained their body temperatures at 35–36°C throughout the experiments. An aperture was made next to the limbus through the sclera with a 30-gauge disposable needle, and a 33-gauge unbeveled blunt-tip needle on a Hamilton syringe was introduced through the scleral opening into the vitreous space for intravitreal injections and introduced through the scleral opening along the scleral interior wall into the subretinal space for subretinal injections. Each eye received 1 μl in intravitreal or 0.4 μl in subretinal injection of vehicle (PBS) or vector at a titer of $1.0 \times 10^{12}$ vg/ml (AAV-2 and AAV-DJ) or $1.0 \times 10^{11}$ vg/ml (AAV-6) .

**Multielectrode array recordings**

All of the procedures were performed under dim red light. The mice were anesthetized and euthanized by quick cervical dislocation. Following enucleation, the retina was dissected at room temperature in Ames' medium bubbled with 95% O2/5% CO2 (A 1420; Merck). The separated retina was placed on a cellulose membrane, and RGC was directed to the electrode and was gently contacted against MEA (MEA2100-Systems; Multi Channel Systems, Reutlingen, Germany) under suction pressure. During the experiment, the retinas were continuously perfused with Ames' medium bubbling at 34 ° C. at a rate of 1-2 ml/min. Recorded signals were collected, amplified, and digitized using MC Rack software (Multi Channel Systems). Retinas were perfused for 30 min in darkness before recording responses. 400, 470, 525, 570, 610, 630 and 660 LED was used in spectral sensitivity examination and white LED was used in the other experiment.

Stimulation was presented for 1 seconds at 60-second intervals. Signals were filtered between 200 Hz (low cutoff) and 20 kHz (high cutoff). A threshold of 40 μV was used to detect action potentials, and action potentials from individual neurons were determined via a standard expectation–maximization algorithm using Off-line Sorter software (Plexon, Dallas, TX, USA). The results were plotted using NeuroExplorer software (Nex Technologies Colorado Springs, CO, USA).

**ERG analyses**

Scotopic ERGs were recorded according to a previous report [1]. Animals were dark-adapted for 12 h and prepared under dim red illumination. The mice were anesthetized with a combination of midazolam, medetomidine and butorphanol tartrate at doses of 4 mg/kg, 0.75 mg/kg and 5 mg/kg of body weight, respectively and were placed on a heating pad that maintained their body temperature at 35–36°C throughout the experiments. The pupils were dilated with a mixed solution of 0.5% tropicamide and 0.5% phenylephrine (Mydrin-P; Santen, Osaka, Japan). The ground electrode was a subcutaneous needle in the tail, and the reference electrode was placed subcutaneously between the eyes. The active contact lens electrodes (Mayo, Inazawa, Japan) were placed on the corneas. Recordings were performed with a PuREC acquisition system (Mayo). Responses were filtered through a bandpass filter ranging from 0.3 to 500 Hz to yield a- and b-waves. White LED light stimulations of 10.0 log cd-s/$m^2$ were delivered via a Hemisphere LS-100 Stimulator (Mayo). The amplitudes were measured and analyzed based on ISCEV (International Society for Clinical Electrophysiology of Vision) standard.

**VEP analyses**

The measuring electrodes were placed more than one week before the measurement. The mice were anesthetized with a combination of midazolam, medetomidine and butorphanol tartrate at doses of 4 mg/kg, 0.75 mg/kg and 5 mg/kg of body weight, respectively. The animals were placed in a stereotaxic holder. A stainless-steel screw (M1.0×6.0 mm) inserted through the skull into the both visual cortex (1.5 mm laterally to the midline, 1.5 mm anterior to the lambda), penetrating the cortex to approximately 1 mm, served as a measuring electrode. Animals were dark-adapted for 12 h and prepared under dim red illumination. At the time of the measurement, the mice were anesthetized again with the same doses. Visual stimuli were generated by a white LED (3 cds/$m^2$ ). Signals were acquired and analyzed with a PuREC acquisition system (Mayo). Signals were low-pass filtered at 300 Hz and averaged over the 60 trials.

**LDT recording**

Mice were tested in a 30 × 45 × 30-cm box, containing equally sized light and dark chambers connected by a 5 × 5-cm opening via which mice could move freely. The bright half of the box was illuminated from above by a white LED. The illumination intensity of measured at the floor level. The animals were placed in the bright half and movement recorded (HD Pro Webcam C920, Logitech, Lausanne, Switzerland). A trial lasted 10 min, and then the testing apparatus was dismantled and cleaned with 70% ethanol. Videos were analyzed using ANY-maze tracking software and were validated by comparison with manual analysis. Time spent in the bright half was recorded.

**VRT recording**

Mice were tested in a 216 × 148 × 220-mm box, containing equally sized light and dark chambers connected by a 120 × 60-mm opening via which mice could move freely. The size of the tablet was 107 × 9.9 × 193-mm (B1-760HD, Acer Inc, New Taipei, Taiwan). The resolution of the display was 1280 × 720 pixels, and the resolution of the videos was 640 × 480 pixels. The luminance of all videos was 20 ± 3 lux. All videos were presented without sound. The box was illuminated from above by a white LED with 10 lux. The illumination intensity of measured at the floor level. The animals were placed in the bright half and movement recorded (HD Pro Webcam C920, Logitech, Lausanne, Switzerland). A trial lasted 15 min, and then the testing apparatus was dismantled and cleaned with 70% ethanol. Videos were analyzed using Move-tr/2D tracking software (Library, Tokyo, Japan) and were validated by comparison with manual analysis. Time spent in the bright half was recorded.

**Gi/o coupled GPCR activation assay**

HTRF-based cAMP detections were conducted with cAMP Gi kit (Cisbio #62AM9PEB, Bedford, MA) according to the manufacturer's instructions. HEK293T cells were kept in DMEM (12-well plate) supplemented with 10% (v/v) fetal bovine serum in a humidified incubator at 37°C 5% CO2. HEK293T cells were seeded on a 12-well plate at $1x10^5$ cells/well, and on the day 2, $1x10^6$ vg/well/500 μl of AAV vector (AAV-DJ-GAGGS, AAV-DJ-CAGGS-GHCR, AAV-DJ-CAGGS-coGHCR) was added and transfected. Transfected cells were kept in the dark for 2 days. On the day 4, after seeding in a 384-well plate at

6,500 cells/well/5 ul and incubating for 4 hours in the dark. Photo-stimulation (525 nm LED $10^{16}$ photons/cm$^2$/s 1 minute) was performed. The signal was detected using plate reader Infinite M1000PRO (Tecan, Männedorf, Switzerland).

**TEM**

Eye cups were fixed with aldehyde/DMSO at 37 °C for 2–4 h and then eye cups were cut in half on their dorsal-ventral axis and fixed again for several minutes. Ultrathin sections were cut with a diamond knife. Specimens were examined using a transmission electron microscope (JEM-1400Plus).

**Western blot analysis**

Isolated mice retinas were homogenized in 80μl of lysis buffer (PBS containing 1% NP-40, 0.5% sodium deoxycholate, 0.1% sodium dodecyl sulfate (SDS), and protease inhibitor cocktail; 1:30, Merck), and centrifuged (15,000g at 4°C for 30 minutes). The supernatant was retained, mixed with an equal amount of sample buffer, and denatured. For gel electrophoresis, 20μg of total protein was loaded onto 12% SDS polyacrylamide gels. The proteins were transferred to nitrocellulose membranes (Pierce Biotechnology, Inc., Rockford, IL) with 30V overnight. The membranes were blocked with 5% skim milk (Bio-

Rad Laboratories, Inc., Hercules, CA) in 1% TBS, followed by incubation with the primary antibodies, anti-CHOP (L63F7) Mouse mAb (#2895, Cell Signaling Technologies, MA), anti-BiP (C50B12) Rabbit mAb (#3177, Cell Signaling Technologies), anti-PERK (C33E10) Rabbit mAb (#3192, Cell Signaling Technologies), anti-ATF-6 (D4Z8V) Rabbit mAb (#65880, Cell Signaling Technologies), anti-pIRE1 Rabbit pAb (ab48187, Abcam), anti-XBP-1s (E8C2Z) Mouse mAb (#27901, Cell Signaling Technologies), anti-ATF-4 (D4B8) Rabbit mAb (#11815, Cell Signaling Technologies) in TBST/blocking solution. Then the membrane was incubated with a horseradish peroxidase–conjugated goat antibodies (1:10000, Zymed) for 2 hours. For the secondary immunoreaction, the PVDF membrane was incubated with WB Stripping Solution (Nacalai Tesque, Kyoto, Japan) to remove antibodies, and blocked again with 5% skim milk in TBS. Further immunoblots were performed using a mouse antibody against β-actin (ACTB, 1:5000, Merck).

**Preparation of cryosections of retinas**

Enucleated eyes were fixed for 20 min in 4% paraformaldehyde (PFA) in PBS and then dissected as previously described [3]. The obtained tissues were post-fixed overnight in 4% PFA in PBS and stored in methanol at –20°C. Cryosections of retinas (12 μm) were prepared as previously described [4], after the eyeballs were immersed overnight in 4% PFA. The retinal sections were observed using a confocal microscope (LSM710; Carl Zeiss, Jena, Germany).

**TUNEL assay**

After the cryosection mentioned above, cell apoptosis was detected by TUNEL using ApopTag In Situ Apoptosis Detection Kits (Chemicon International, Darmstadt, Germany; cat. #S7165) according to the manufacturer's instructions. Nuclei were counterstained with DAPI. The retinal sections were observed using a confocal microscope (LSM710; Carl Zeiss, Jena, Germany).

**OCT Imaging**

The thickness of the retina was analyzed by an SD-OCT system (Envisu R4310; Leica, Wetzlar, Germany) tuned for mice. The imaging protocol entailed a 3 mm×3 mm perimeter square scan sequence producing a single en-face image of the retina through a 50-degrees field of view from the mouse lens, following mydriasis. The en-face image consisted of 100 B-scan tomograms with each B-scan consisting of 1000 A-scans. The retinal thickness of 150 μm from the optic disc of each quadrant was measured.

**Quantification and Statistical Analysis**

All of the results are expressed as the mean ± SEM. The averaged variables were compared using the unpaired t-test and the one-way ANOVA test. P-values of less than 0.05 were considered statistically significant. All experiments were randomized.

**HPLC analysis of Retinal**

After 15 hours of dark adaptation, mice were exposed to light adaptation at 1000 lux for 10 minutes. The mice were subsequently sacrificed, and the removed mouse retinas were homogenized. Hydroxylamine was added to the homogenized retinas for oximation. The retinal- oximes was dissolved in hexane to make a sample for HPLC analysis. All of these processes were performed under dim red lights. Two retinas were used per assay.

Retinal oximes were analyzed by HPLC (Shimadzu LC20A series, Japan) with a silica column (Ultrasphere 5um, SI 250 x 4.6mm, Avantor, USA). The mobile phase consisted 96.0%(v/v) hexane, 4.0%(v/v) ethyl acetate and the flow rate was 1.0mL/min. The column temperature was 35°C. Absorbance at 360 nm was monitored for retinal oximes. Each retinal isomer was quantified from the area of the corresponding peak based on a calibration retinal standard reagent. All trans-retinal (Sigma-Aldrich) and 11-cis retinal (Toronto research chemicals) were used as standard reagents.

### Data and Software Availability

Raw MEA spike data were sorted offline to identify single units using Offline Sorter software (version 4.4.0)(Plexon). Spike-sorted data were analyzed with NeuroExplorer 5 software (version 5.115) (Nex Technologies). The data that support the findings of this study are available from the corresponding author on request.

### Supplemental Figures

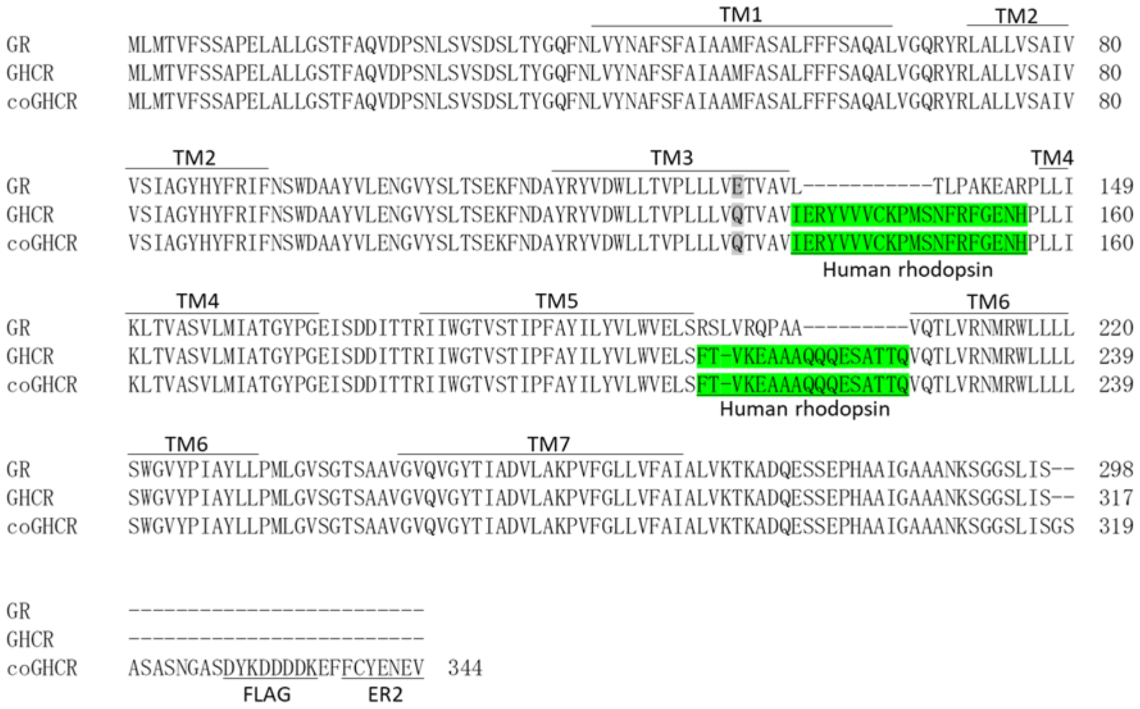


**Figure S1. GHCR structural alignment**

Primary structure alignment of wild type *Gloeobacter* rhodopsin (GR), GHCR, and coGHCR. Gray, E132Q mutation; green, inserted constructs of the second and third loops of human rhodopsin. TM, transmembrane domains; FLAG, FLAG tag; ER2, ER2 ER export signal.

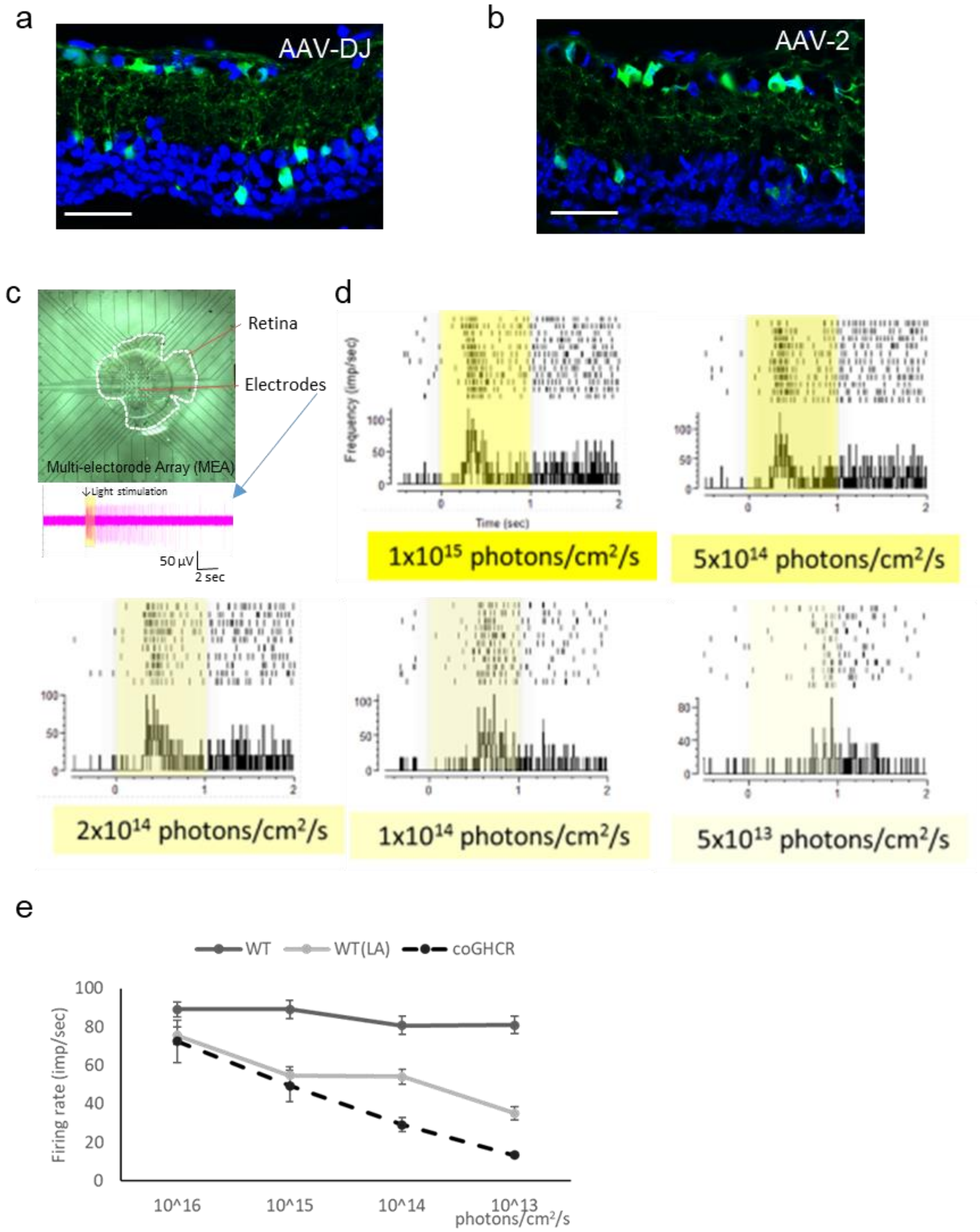


**Figure S2. Ectopic expression of AAV-2-CAGGS-GHCR restores light responses in the *rd1* mouse retina**

(a, b) Confocal images of transverse sections of a C57BL/6 retina 2 months after AAV-DJ-CAGGS-EGFP (a) and AAV-2-CAGGS-EGFP (b) intravitreal injection. Blue, DAPI nuclear counterstaining. Scale bar, 50 μm. (c) MEA schematic. MEA is used to measure the extracellular potential of RCGs in contact with the electrode ex vivo. (d) Raster plots and peri-stimulus time histogram of light stimulation of GHCR-treated (AAV-2-CAGGS-GHCR) mice. The response to stimulation with a white LED with varying light intensity (irradiation duration, 1.0 s) was recorded. (e) Quantification of firing rates from RGCs in wild-type mice, light-adapted (LA) wild-type mice and rd1 mouse retina 2 after AAV-DJ-CAGGS-coGHCR intravitreal injection at the indicated light intensity.

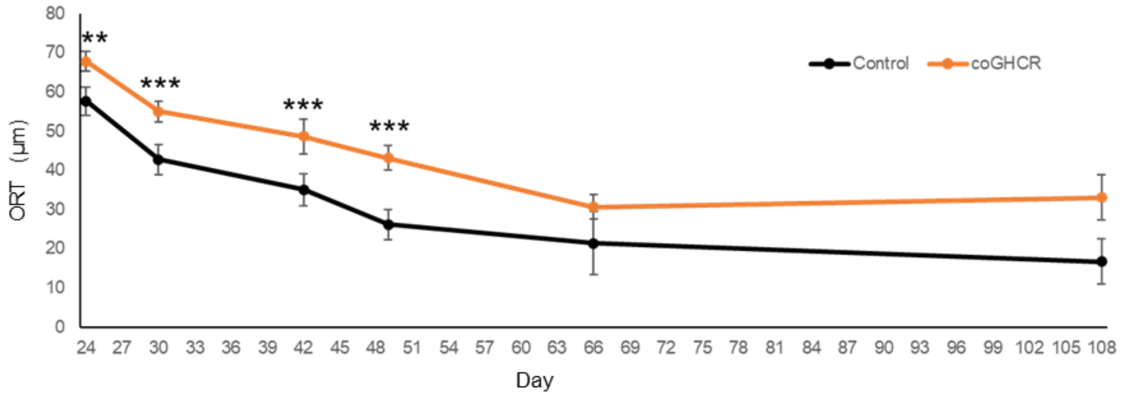


**Figure S3. Characterization of coGHCR-mediated retinal protection via OCT imaging**

Follow-up of ORT (from ONL to cone outer segment) after AAV-DJ-CAGGS-coGHCR and control (AAV-DJ-CAGGS-EGFP) subretinal injection, measured with OCT. Error bars represent SEM. Data were analyzed with the unpaired t-test; ** represents $p < 0.01$ and *** represents $p < 0.001$.

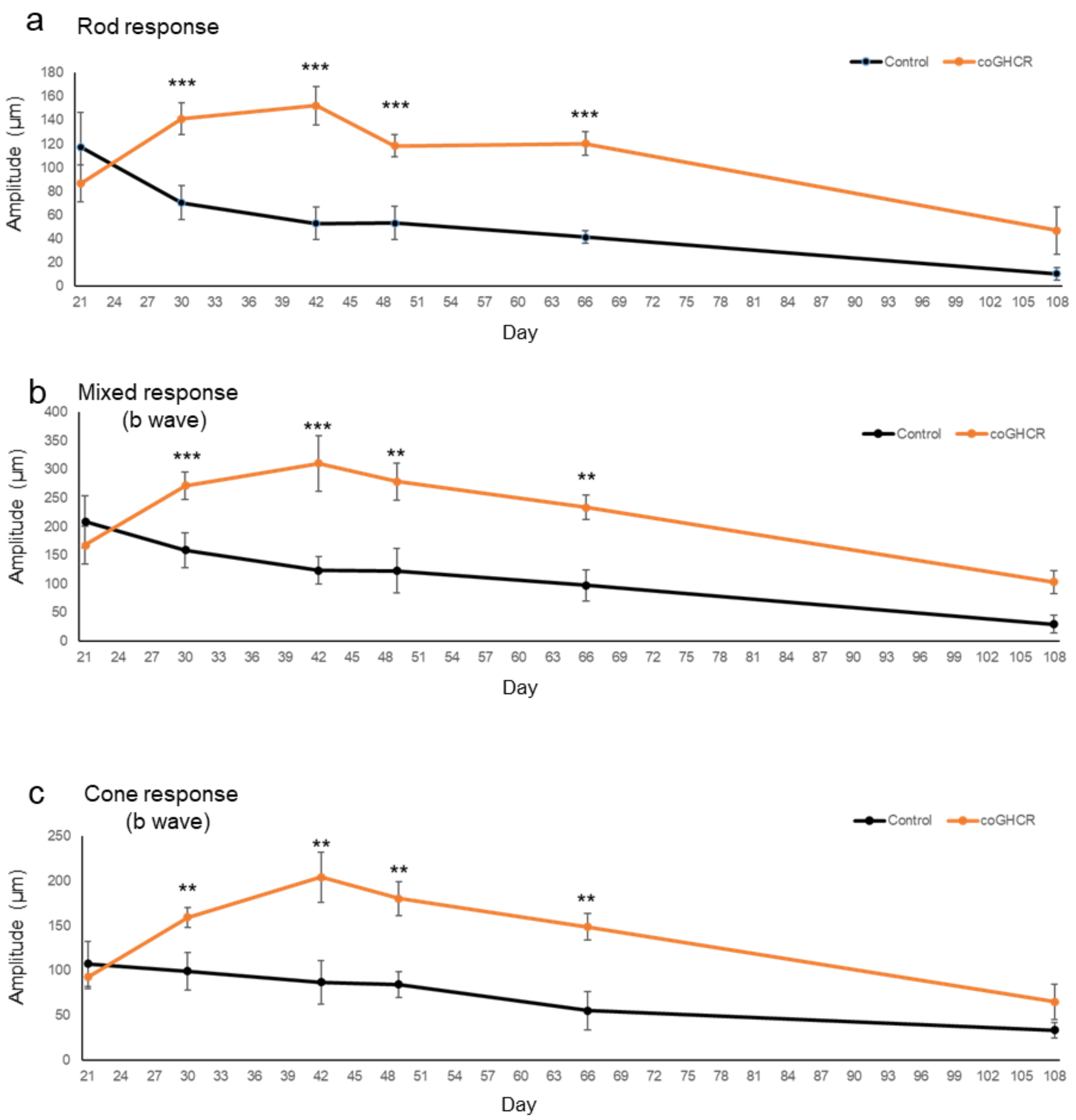


**Figure S4. Characterization of coGHCR-mediated retinal protection as assessed by ERG recording**

(a–c) Follow-up of the ERG amplitude of the rod response (a), the mixed b wave (b), and the cone b wave (c) after AAV-DJ-CAGGS-coGHCR and control (AAV-DJ-CAGGS-EGFP) subretinal injection. Error bars represent SEM. Data were analyzed with the unpaired t-test; ** represents $p < 0.01$ and *** represents $p < 0.001$.

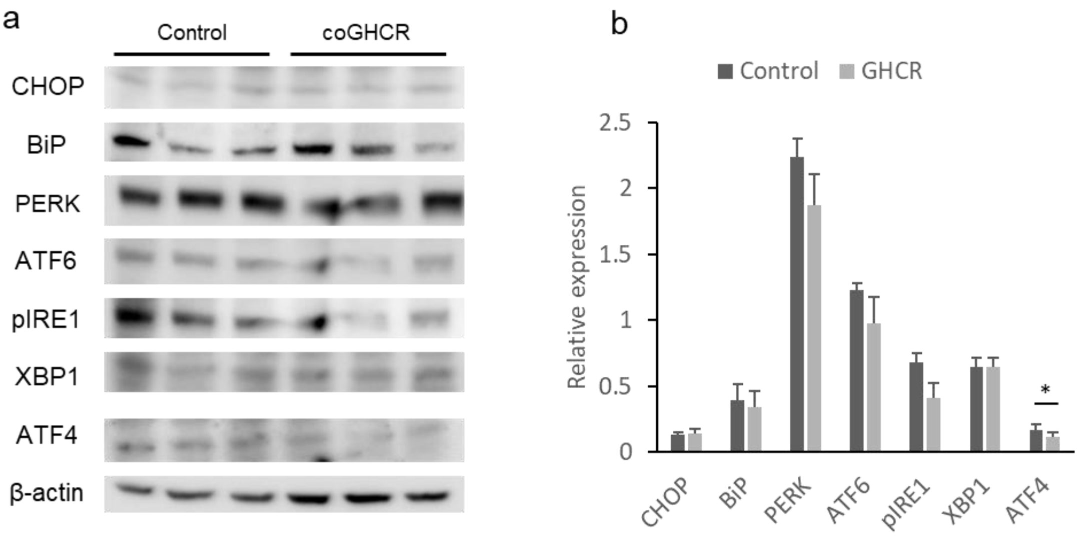


**Figure S5. Expression of ER stress marker proteins in coGHCR-treated and control mice at PND 31**

(a) Expression of ER stress marker genes in coGHCR-treated and control mice at PND 31.

(b) Quantitative analysis of the expression in relationship to β-actin expression, as an internal control. Error bars represent SEM. Data were analyzed with the unpaired t-test; * represents $p < 0.05$.
